# BioBloom, a method for barcoded saturation mutagenesis of an entire bacterial genome

**DOI:** 10.64898/2026.02.04.703572

**Authors:** Max Schubert, Edward Sukarto, Fatima R. Martin, Jordan E. Mancuso, Amy Spens, Tom Pedersen, Keren Isaev, Harley Greene, Nathan D. Hicks, Una Nattermann, Jonathan Liu, Devon A. Stork, Erika A. DeBenedictis

## Abstract

Saturation mutagenesis is a powerful tool for understanding and engineering the function of biological systems. and has been applied successfully to characterize the mutational landscape of individual proteins and genetic loci. However. it has not been applied at the whole-genome scale due to the challenges of both creating and quantifying a saturating set of mutations. Here we introduce BioBloom. a retron-based method for barcoded saturation mutagenesis at the scale of a whole bacterial genome. We constructed a barcoded BioBloom library with >99% projected sampling of saturating single - nucleotide polymorphism (SNP) mutations of the *E. coli* genome, and applied it to identify beneficial mutations under salt and antibiotic selection. Relative to other techniques like CRISPR-enabled mutagenesis or Adaptive Laboratory Evolution, BioBloom excels at identifying diverse causal SNPs quickly and at smaller working volumes. A barcoded, saturating mutation library is also a shared resource, and we are releasing the updated BioBloom-Ec.-2.0 library to the scientific community for broader adoption and application. Together. BioBloom makes barcoded saturation mutagenesis accessible at whole-genome scale, creating new opportunities for large-scale data collection and bacterial engineering.

## INTRODUCTION

A fundamental goal of modern biology is to understand the relationship between genotype and phenotype. Pooled assays have revolutionized our ability to generate genetic variation and observe its effect at scale, enabled by advances in DNA synthesis and sequencing. For example, saturation mutagenesis involves creating and measuring the complete set of nucleotide-level changes for single proteins or loci^1^. By “saturating” all possible mutations, we learn about the role each nucleotide plays, yielding insights into protein folding, function. domains, active sites, and strategies for engineering^2-4.^ In contrast, TnSeq or CRISPRi are “barcoded knock-out” assays, efficiently querying loss-of-function mutations across all genes in a genome, but do not provide the nucleotide-level resolution of saturation mutagenesis. The value of both types of large, high-quality genotype-to-phenotype datasets is only enhanced by advances in Al, where such datasets can be used to train predictive^5^ and validate generative^6^ models of biology.

However, to date there are no pooled screening methods that enable saturating mutagenesis across entire microbial genomes. There are a number of methods for genome-wide loss-of-function screens, notably TnSeq. These methods have yielded invaluable insight about gene function and are widely used by the field^7–9^, but create “knock-outs” lacking nucleotide resolution and destroying rather than altering gene function or expression. Small mutations including SNPs do explore these avenues, and beneficial mutations are commonly identified through Adaptive Laboratory Evolution (ALE). ALE is well suited to identify Single Nucleotide Polymorphisms (SNPs) that improve performance across industrial^10,11^ and research applications^12^ and is employed as a strain engineering technique.

However, ALE can take hundreds of days in total, especially when growth is slow^12,13^. In addition, determining which specific ALE mutations provide benefit requires separate follow-up experiments^14,15^ which are often labor-intensive and low-throughput^16,17^.Thus while ALE can often generate improved strains, it is unable to generate data at the scale of whole-microbe-genome saturation. New techniques for microbial whole-genome saturation mutagenesis have the potential to overcome the limitations of these approaches, resulting in scientific insight, rapid strain engineering, and valuable datasets for AI. These methods could achieve the genome-wide breadth and data generation of methods like TnSeq, while exploring the effects of small mutations, which have been shown to be useful and illuminating in saturation mutagenesis and ALE.

Here, we report BioBloom, a pooled method for barcoded, genome-wide site-saturation mutagenesis, and its implementation in *E. coli*. This method uses retron recombineering to create a genome-wide saturating set of SNPs, while linking an identifying sequence to each mutant for tracking across the entire mutant pool. This pilot study explores nearly all approximately 15 million Single-Nucleotide Polymorphism (SNP) variants along with wild type sequences for the entire *Escherichia* coli MG1655 genome. We first demonstrate construction of this library, sampling a projected 99% of all saturating SNPs. We then apply this method to test the functional contribution of every SNP to salt and antibiotic tolerance. Following selection on antibiotic, we detect approximately 26% of expected beneficial SNPs, as well as some novel SNPs. In a salt tolerance experiment, we detect hundreds of beneficial SNPs, finding that beneficial SNPs cluster within biologically meaningful regions such as genes, regulatory regions, and specific protein catalytic sites and domains. Together, these advances position BioBloom as a scalable. high-resolution platform for pooled genome-wide mutation screening and data collection in bacteria.

## RESULTS

### Barcoded Mutant Libraries Through Retron Recombineering

Previous work has used retron recombineering to conduct pooled screening of mutations in bacteria^18^. Like CRISPR, retrons originally evolved as components of bacterial immune systems^19^; part of this function requires producing single-stranded DNA through targeted reverse-transcription^20^. In retron recombineering, a retron containing a 70-100 bp “pro-donor” sequence bearing a mutation of interest is introduced to the cell on a plasmid (Figure 1a). Once induced, the retron uses this sequence as a template to generate single-stranded DNA donors. When co-expressing a suitable Single-Stranded Annealing Protein (SSAP), this donor is directed to the replication fork where it anneals to edit the genome^**1**8^,^**2**1^.^**22**^ (Figure 1a). The process of propagating the mutation to the genome is efficient: the edited fraction of cells is approximately 50% or higher across a range of loci after reasonable induction time^18^. and longer induction time^18^ or new approaches like alternative or engineered retrons can further improve editing^23–25^. Retron recombineering results in a specific mutation of the bacteria’s genomic DNA, while the pro-donor sequence specifying this mutation is retained *in trans* on a plasmid. The retained pro-donor sequence can double as an amplicon-sequencing barcode, enabling pooled quantification of genomic mutant abundance and fitness by Next Generation Sequencing (NGS)^18^.

Using retron recombineering for genome-scale saturation $NP mutagenesis requires creating a retron plasmid library containing one pro-donor sequence for every possible SNP. For a bacterial genome of a given length, there exist 3 alternative bases at each position. For the E. *coli* MG1655 genome (4.6 × 10^6^ bp) this corresponds to roughly 1.4 × 10^7^ SNPs, which could be specified by synthesis (Figure Slb). Synthesis of > 10^7^ oligos of sufficient length would cost ~$100k at this time, despite historically low oligonucleotide pool synthesis cost^26^.One simple cost reduction strategy is to use degenerate “N” nucleotides rather than explicitly specifying each SNP. cutting synthesis required by a factor of three. We thus designed a donor sequence for each base in the genome. with a degenerate “N” mixture specified at a central editing nucleotide (Methods) and purchased pooled synthesis of these 4.6 × 10^6^ oligos for less than $30k. The resulting pool contains approximately 1.8 × 10^7^ sequences including all SNPs as well as wildtype at each position. This library is about three orders of magnitude larger than the largest retron libraries reported to date^27^, but larger libraries are created for other applications^28,29^,suggesting that it should be feasible to construct. The oligo pool was amplified by PCR, assembled into a vector using a Golden Gate approach, and transformed, achieving 2.8 × 10^9^ CFU. or > 100-fold representation of our library (Methods, Figure 1b). This plasmid library was then transformed into an E. *coli* MG1655 background optimized for retron editing^18,30^achieving even higher coverage of >1 × 10^10^ CFU. Retron recombineering was then induced in triplicate cultures to produce whole-genome saturation library BioBloom-Ec.-1.0 (see Methods. Table Sl).

**Figure 1.**
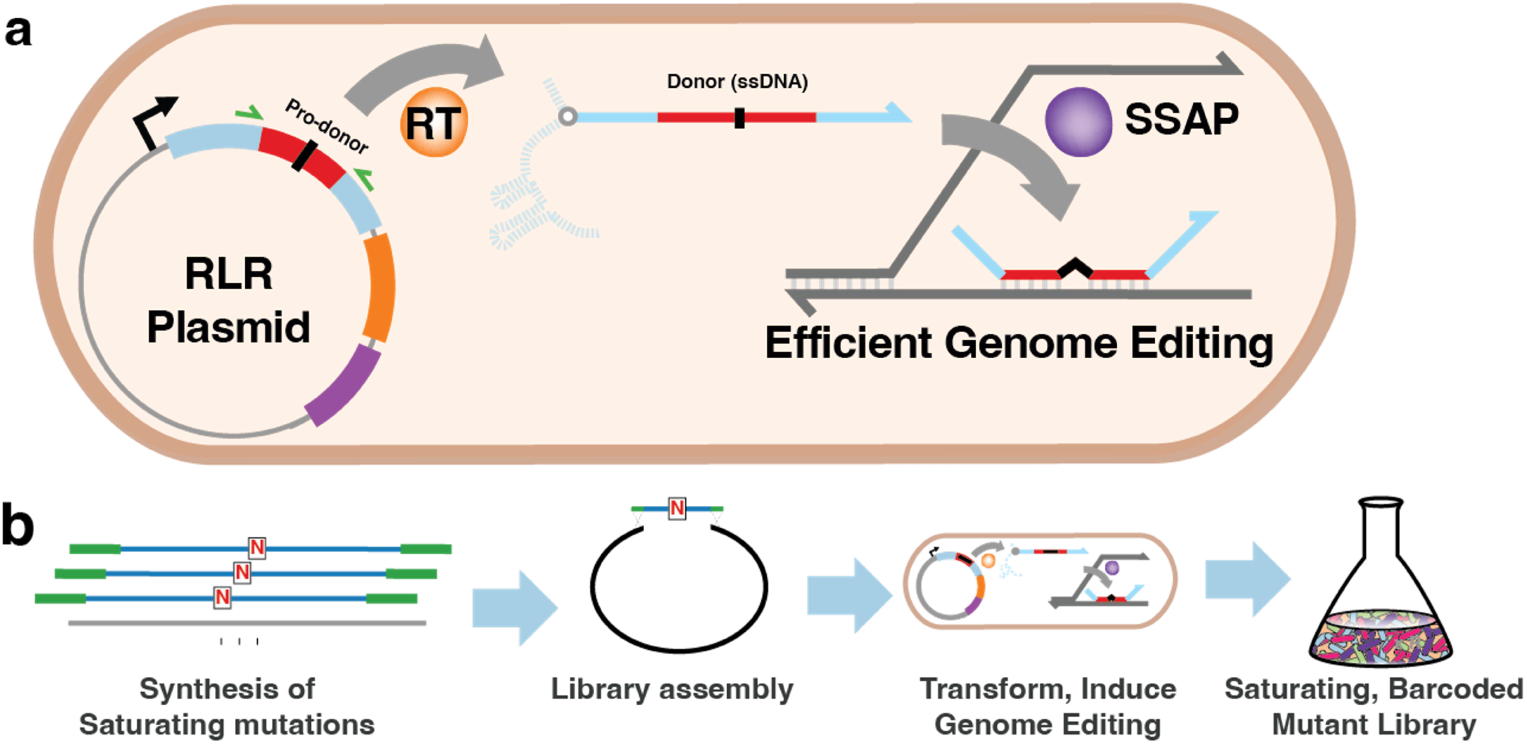
Retron recombineering and creation of BioBloom libraries. **a)** Retron Library Recombineering (RLR) can be used to make genomic edits while retaining a unique barcode for each variant. The Retron Plasmid encodes a Pro-donor sequence, retron Reverse Transcriptase (RT, orange). and a single-stranded annealing protein (SSAP, purple). The Pro-donor sequence is transcribed and targeted by RT, producing a Donor single-stranded DNA (ssDNA) including single nucleotide polymorphisms (SNPs) and surrounding homology to the genome (red). This donor is recruited to the replication fork where it creates precise edits, catalyzed by Single-Stranded Annealing Protein (SSAP. purple) expressed from the Retron Plasmid. The Pro-donor is retained on the plasmid after the edit, where it is available for amplicon-sequencing using specific primers (green), acting as a barcode to identify the mutant. **b)** Donor sequences 82bp were designed for each genomic position containing a degenerate “N” at the central base. These sequences were synthesized with flanking golden gate adaptors (green). assembled into a plasmid library, then transformed and induced in recipient cells to create the barcoded BioBloom mutant library.

### Saturation Analysis

Having constructed retron libraries, we next sought to estimate the fraction of barcoded. saturating SNPs contained in these libraries. Using a coupon collector model (Methods), we can estimate the fraction of all ~1.5 × 10^7^ SNPs sampled as the total library CFU achieved when constructing the library increases. Assuming random sampling with replacement from a uniform pool, and accounting for the 25% of our library members which encode wild-1ype sequences, the 2.8 × 10^9^ CFU library we constructed is expected to sample >99% of all SNPs (Figure 2a). In realily, the frequency distribution of library members is unlikely to be uniform, with pooled libraries tending to follow a log-normal frequency distribution^31–32^. We thus expanded our model to consider sampling from pools with log-normal spread or skew summarized by***σ***, the standard deviation after log-transformation. For example with ***σ=*** 1, the most frequent 10% of donors claim ~35-40% of all observations, and with ***σ***= 2. the most frequent 10% claim ~75-80% of all observations. Nonetheless, even with a substantially skewed library having ***σ***= 2, we would expect to sample more than 90% of SNPs given the library size obtained (Figure 2a). In order to estimate the skew and saturation of our final edited library we had constructed, we sequenced pro-donors of the edited mutant library with ~10^8^ reads for each replicate, fit a log-normal distribution to the coverage across genome positions, and found ***σ***is approximately 1.3 for all three replicates after induction and editing (Figure S2a). This indicates our library is expected to sample more than 99% of all possible single f. *coli* SNPs.

**Figure 2.**
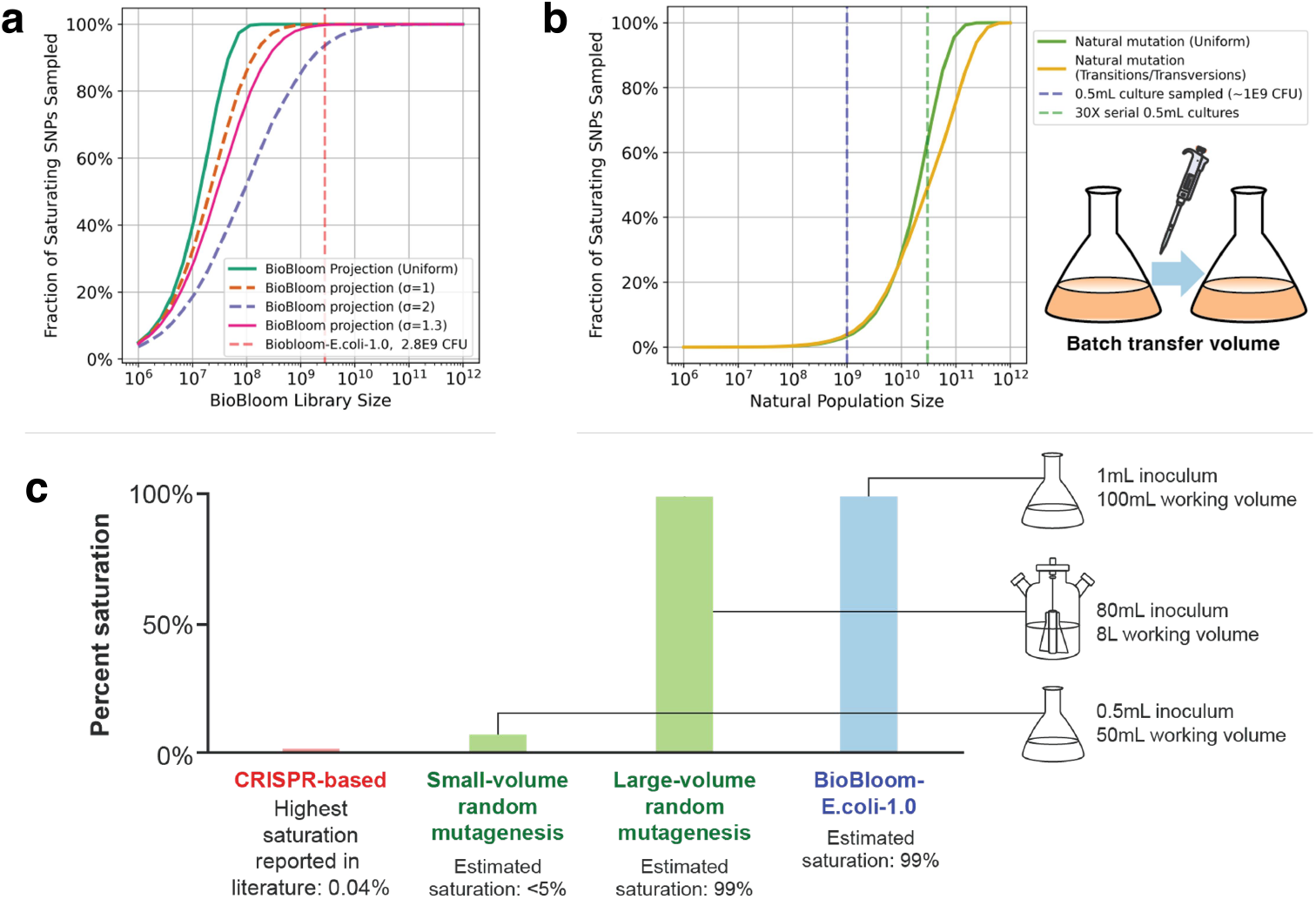
Analysis of library saturation. **a)** Model of constructing a saturating mutagenesis library. The traction of all saturating SNPs obtained (y axis) is shown as a function of library size (x axis). A coupon collector model is used to estimate the traction of the complete set obtained, when sampling from either a uniform set (green) or with differing estimates of library skew (orange, purple. pink). The estimated BioBloom library size achieved in this work is indicated (dashed vertical line, pink). **b)** Model of saturating SNP sampling. in a natural population. The predicted fraction of all saturating SNPs (y-axis) occurring in a natural population of a given size (x axis). Mutation number is derived by multiplying mutation rate, genome size, and number of divisions given the population size, then subjected to the same coupon collector model as in (a). A uniform mutation rate is modeled (green), as well as a simplified model of unequal mutation rate between transitions and transversions (yellow). Population size tor both a 0.5 ml confluent culture (dotted blue) and cumulative mutations after 30 such cultures in series (dotted green) are indicated to model common situations in Adaptive Laboratory Evolution. **c)** Comparison of the potential genome-wide saturation fraction of BioBloom, CRISPR-based screens, and random mutagenesis and the relationship with the inoculation and working volumes.

Next, we sought to determine the population size, and thus volume of culture, needed to faithfully and practically carry this diverse set of barcoded SNPs into selection experiments. Using the ***σ*** value 1.3 as determined above and factoring in that retron editing does not occur at 100% efficiency, we can adapt our model above to ensure we are passaging enough cells into selection. We determined that l ml of saturated culture was enough to carry >99% of possible SNPs into selection if retron editing is 50% efficient, which has been achieved by similar protocols in other studies^1,6–8^. Even if editing is only 10% efficient, >90% of all SNPs should be sampled during selection (Figure S2b). When conducting selection experiments, it is 1ypical to propagate libraries in enough medium to enable at least 100-fold expansion of the population. Therefore. our l ml inoculum starting cultures would best be selected in 100 ml of working volume of medium. This may not be practical for all studies, and indeed the approximately 60% saturation achieved by a 0.1 ml inoculum volume and 10 ml working volume may be a better fit for some workflows (Figure S2b).

It is worth comparing the fraction of saturation SNPs sampled by BioBloom to that occurring naturally in a population of bacteria. These natural, non-bar coded mutations are the basis of diversification for Adaptive Laboratory Evolution (ALE). It is sometimes assumed that a bacterial culture contains a substantial fraction of all saturating SNPs, but this is generally not true. Existing models estimate that a 5 ml *E. coli* culture would contain 1 × 10^7^ mutations^33^, which is fewer than the −1.5 × 10^7^ saturating SNPs possible, and expected to sample about 49% of mutations given a simple coupon collector model. In practice the fraction sampled is far lower, as small fractions of cultures used to inoculate batch cultures or selection experiments, often at about 1/100th of the final population size such that selection can occur This transfer volume contains fewer cells and thus samples fewer mutations. Adapting our coupon collector model from above, we find that with normal mutation rate, a practical inoculum volume of 0.5 ml is expected to sample <5% of all SNPs (Fig 2b). Serial propagation over 30 subcultures is expected to sample ~65% of saturating SNPs, or <45% when considering the lower rate of transversion mutations (Fig 2c, Methods). Biobloom achieves ~99% SNP saturation with a 1 ml inoculum and 100 ml working volume, and doing the same with natural populations would require an 82 ml inoculum and a 8 liter working volume. Increasing mutation rate can sample more SNPs, but results in strains with far more mutations per genome that are difficult to deconvolute, which we discuss in the next section. Thus, we expect Biobloom to sample a far larger fraction of saturating mutations in volumes that are practical for scaled, replicated experiments. Combined with the benefits of amp icon sequencing for efficiently, precisely quantifying mutation frequency, we predict BioBloom will make saturating SNP studies far more practical.

The Biobloom method is also expected to have advantages over CRISPR based barcoded mutagenesis a pp roaches. A number of recent studies have identified methods that allow precise CRISPR editing in microbes, including lnscripta −Onyx34, CREATE^35^, MAGESTIC^36^, and CRISPEY^37^. Biobloom libraries vastly exceed the size and saturation percentage of the largest mutagenesis libraries created by these CR ISPR based approaches to date, all of which are 50, 000 members or smaller, able to sample less than 0.4% of all SNPs in the respective microbial genomes (Table S2, Figure 2c). CRISPR-based app roaches ca n only introduce mutations near PAM sites in the genome, limiting their estimated theoretical maximum to 75% saturation editing of the *E. coli* genome^35^. Additional practicalities are in favor of retron based method when very large libraries are desired, including : 1) CR ISPR based methods require both a guide RNA and a repair template, complicating library construction; 2) the synthesized regions are lengthy and thus more costly requiring 200bp^36^ or 250bp^35^ of total synthesis, approximately double the length used in this study and 3) double-stranded breaks are toxic to microbial cells, making construction of a large even library challenging. Together although not impossible in theory CRISPR-based mutagenesis is a less promising approach for generating microbial whole-genome saturation libraries relative to retron mutagenesis, and unsurprisingly the Biobloom library we created exceeds the size of any reported microbial CRISPR-based library by more than two orders of magnitude

### Validation Of Saturation Library With Selection On Antibiotic

Having created the whole-genome saturation library Biobloom-Ec 1.0 we next sought to validate it by exploring known antibiotic resistance mechanisms. Mutations that confer rifampicin resistance are well characterized, with the majority of high - resistance mutations occur ring in *rpoB* the gene encoding the protein target of rifampicin Diverse rpoB mutations lead to resistance at least 65 amino acid changes encoded by 69 SNPs^38^ have been reported These mutations include al 12 possible single-base transition/transversion mutations^39^. While resistance mutations exist outside of rpoB these tend to confer more modest resistance and are not sufficient to tolerate high rifampicin concentrations^40^. Together, selection on rifampicin presents a first simple test case with a large number of known ‘solutions’ that should be recoverable from a whole-genome saturation library.

We subjected Biobloom-Ec.-1.0 to selection on rifampicin in triplicate and sequenced donor/barcode amplicons after selection (Figure 3a). Mutant alleles were identified by alignment to the genome, and a permutation test was used to quantify how unlikely each observed allele frequency is under the empirical data distribution; these p-values were then combined across replicates, and p-values were adjusted using Benjamini-Hochberg correction (see Methods). The *rpoB* locus, where the majority of high-fitness mutations are expected, gives rise to the most significant SNPs (Figure 3b). 18 SNPs are detected at the *rpoB* locus with adjusted p< 0.05, or 38 SNPs with p< 0.05 before adjustment, clustering at sites of known rifampicin resistance mutations (Figure 3c). About half of these 38 mutations have been previously described, representing about one-quarter of known rifampicin resistance mutations (Figure S3c). Four SNPs representing known *rpoB* alleles V146F, D516 Y, D516V and I 572 F were observed in ~1000x g reater abundance than others, likely consuming a majority of reads and restricting our ability to detect a wider set of beneficial alleles; we comment further on how to overcome these limitations in the Discussion.

**Figure 3.**
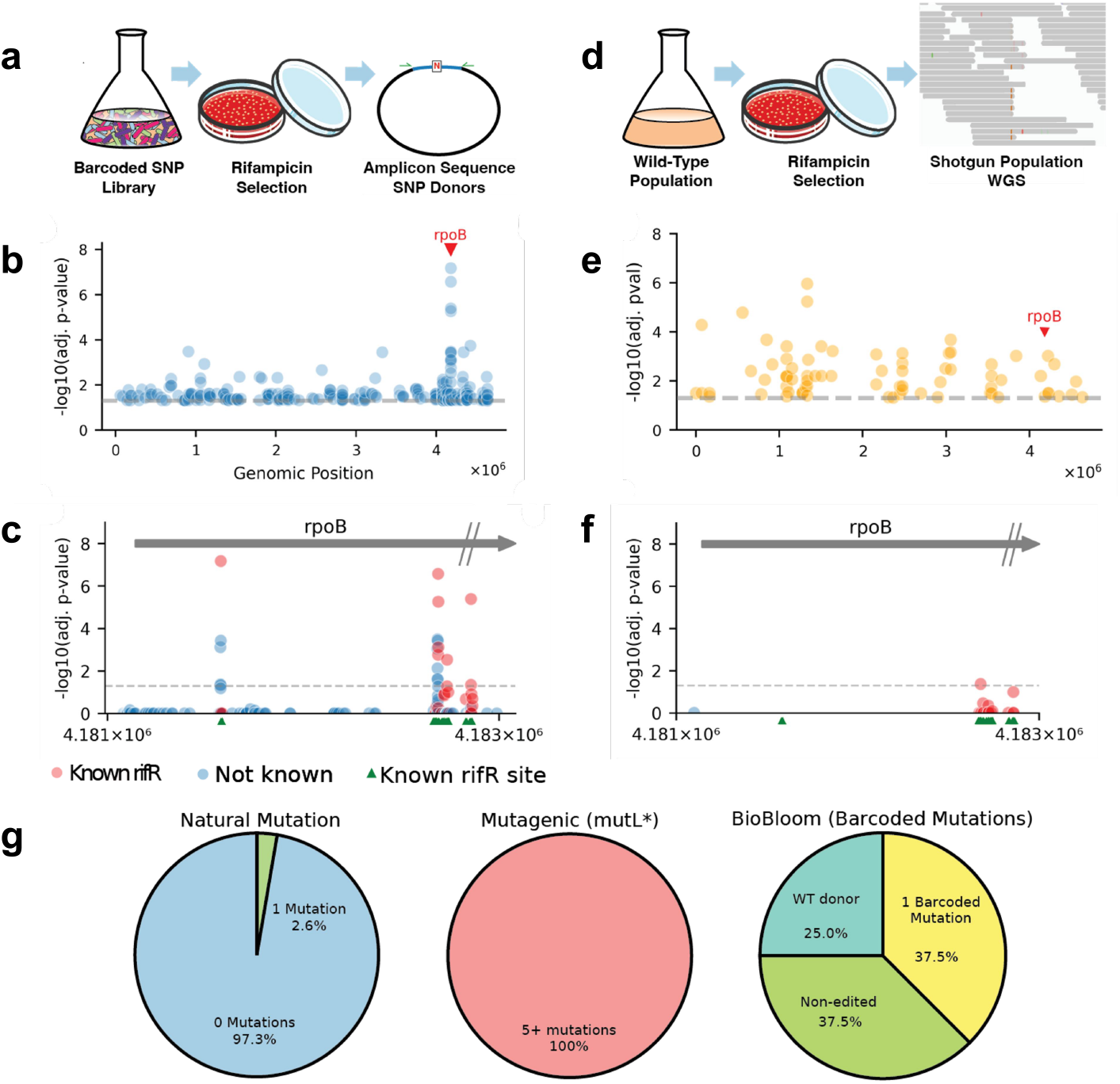
BioBloom applied to antibiotic resistance. **a)** Schematic of a BioBloom rifampicin resistance experiment. A barcoded mutant library is plated on solid rifampicin medium, and then assessed by targeted amplicon sequencing. **b)** Genome-wide results of BioBloom rifampicin resistance experiment. The *rpoB* locus is indicated (red). SNPs with adjusted p-value < 0.05 are depicted at their genome position and the negative log of their adjusted p-value. Adjusted p = 0.05 is indicated with a dashed grey line. c) Results of the BioBloom rifampicin resistance experiment, at the rpoB locus. All mutations detected with 10 or more sequencing counts are shown, adjusted p-value of 0.05 is indicated with a dashed grey line. Known resistance mutations from our literature set are colored red, the remainder are blue. Positions where rifampicin resistance mutations are described in the literature set are depicted in green along the x-axis. The *rpoB* gene sequence is indicated with a grey arrow, with hash marks indicating that the full gene extends beyond the area of the plot. **d)** Schematic of rifampicin resistance experiment using natural or mutagenized populations. Populations are plated on solid rifampicin medium, then assessed by whole-genome shotgun sequencing. **e)** Genome-wide results of natural population rifampicin resistance experiment, depicted in the same manner as in (b). f) Results of the natural population rifampicin resistance experiment at the rpoB locus, depicted in the same manner as in (c). g) Comparison of per-cell mutation count between natural populations, mutagenic populations, and barcoded BioBloom populations. For BioBloom, average editing efficiency is estimated at 50% following literature^19,24^.

We also compared this treatment of Biobloom-Ec. −1.0 to a m ore traditional control reminiscent of ALE, in which cultures of *E. coli*, either experiencing baseline mutation or increased mutagenesis from an induced mutagenesis plasmid^41^, were also selected on rifampicin in quadruplicate using the same culture volumes as the Biobloom experiment (Figure 3d). We used whole-genome population-level shotgun sequencing and low-frequency variant-calling to identify SNPs post-selection. When subjecting the observation counts for SNPs obtained in this way to the same statistical framework as above, we see that SNPs in *rpoB* are not more significant/ abundant than SNPs in other loci genome-wide (Figure 3e). Thus unlike the Biobloom method, this approach would be unable to identify *rpoB* as the primary causal locus without prior knowledge or followup validation. Only 1 SNP within the *rpoB* locus achieved adjusted p< 0 05 in the baseline condition (Figure 3e), or 4 SNPs in the increased mutagenesis condition (Figure S3a, S3 b).

There are likely three major factors at play that cause this traditional mutagenesis to be less effective than Biobloom in identifying causal SNPs. First as discussed earlier natural populations simply contain fewer mutations. With norma mutation rates, only about 2.6% of cells are expected to have 1 mutation, in contrast with the 37 5% we expect with our Biobloom libraries given that 75% of strains obtain non-wild-type donors and we estimate 50% editing in our process (Figure 31) This means that more mutations are sampled, more times, by more cells. Second, in natural populations causal *rpoB* mutations co-occur with unrelated “hitchhiker” or background mutations, indistinguishable in the resulting data. This problem is only exacerbated by applying mutagenesis, which increases the number of mutations sampled, but also accrues multiple mutations per cell (Figure 31) exacerbating the issue of background mutation In contrast Biobloom ‘s barcoded mutagenesis creates and interrogates single mutations specifically and the effect of background mutations is mitigated across greater library coverage or replicates Third, Biobloom ‘s barcoded nature makes it substantially more sequencing-efficient than an approach requiring whole-genome sequencing. “Shotgun” reads of a genome having a single mutation are more than 99% wild-type with about 3 × 10^6^ 150bp reads required to cover a 4.6 ×10^6^ bp genome and observe a mutation-in contrast with the single barcode read required to sample a Biobloom mutant Together these factors highlight the value of barcoded mutagenesis techniques like Biobloom in rapidly identifying causa loc and mutations

### Application Of BioBloom To Salt Tolerance In E. Coli

Next, we applied BioBloom to the more open-ended task of enhancing salt-tolerance in E. coli, which is expected to surface a variely of mutations throughout the genome^42^.^43^. We selected BioBloom-Ec.-1.0 in 5% salt medium in triplicate, conducting three sequential 1/100 dilutions into this medium and growth to saturation, then quantified pro-donor frequency across these cultures using amplicon sequencing (Figure 4a. Methods). After two selection cultures, 1429 SNPs were found with p<0.01 (permutation test) indicating enrichment, with 129 such mutations having p<0.05 after correcting for multiple hypothesis testing. highlighting loci where beneficial mutations can be detected with high confidence (Figure 4b). Notably, after three batch cultures in salt selection, the results are similar but only 785 mutations have p<0.01 before correction, and 42 have p<0.05 after correction (Fig. S4). indicating that earlier timepoints are optimal for obtaining informative data under these assay conditions.

**Figure 4:**
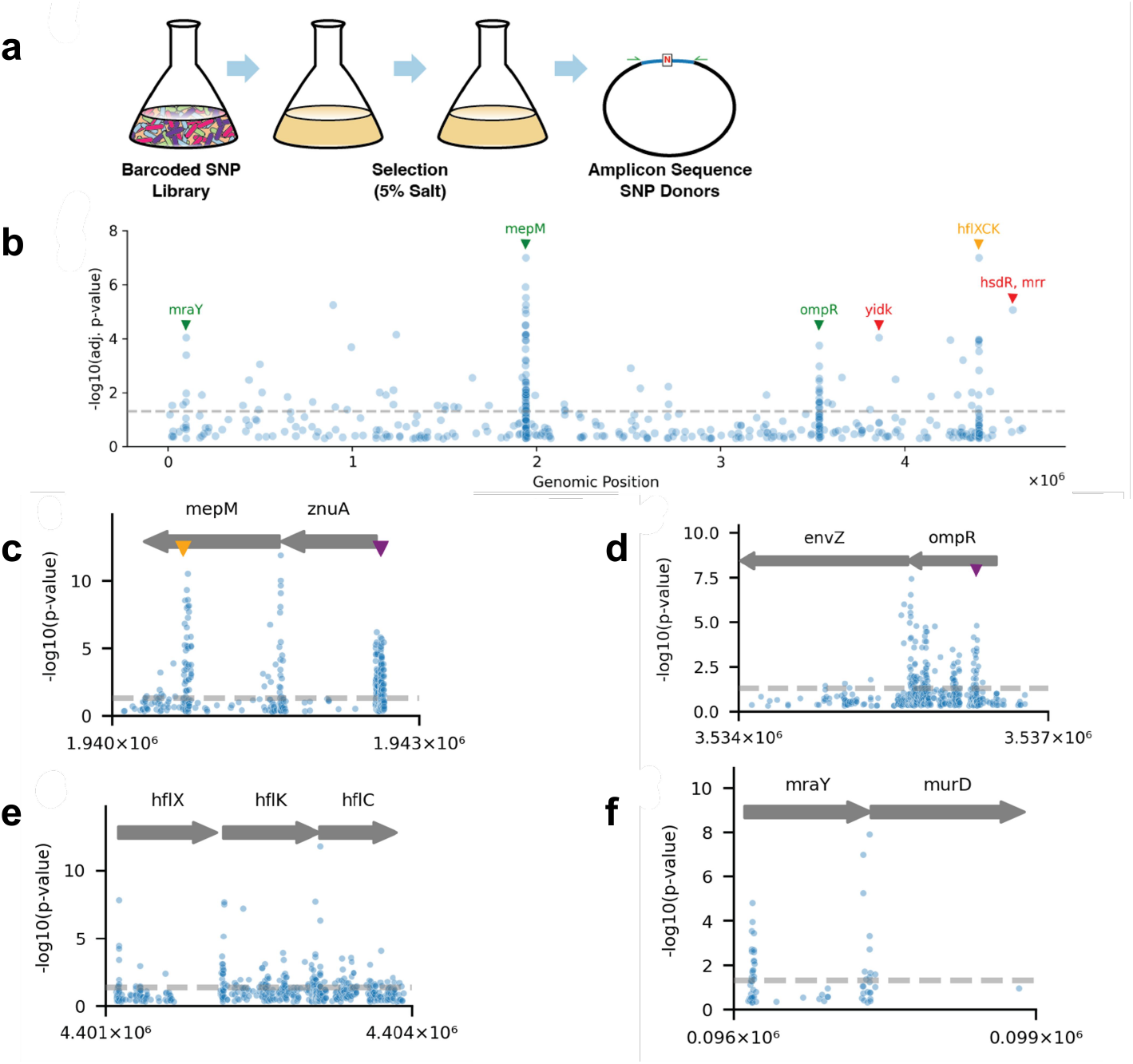
Selection under salt stress. **a)** Schematic of BioBloom experiment investigating salt tolerance. The BioBloom library is cultured twice in 5% salt medium. then assessed by targeted amplicon sequencing. **b)** Genome-wide results of salt tolerance experiment. Significantly-enriched mutations are depicted according to their position in the reference genome (x-axis) and the −log of their adjusted p-value (y-axis). Mutations with adjusted p<0.5 are depicted. with adjusted p = 0.05 indicated with a dashed grey line. *mepM, ompR*. and mraY loci, which we considered plausibly linked to salt tolerance, are indicated in green. The *hflCKC* locus is indicated in yellow. these significant mutations were suspected to be some kind of artifact. Significant *yidK* and *hsd*, mrr alleles are indicated in red, these solitary mutations were expected to be experimental noise. These color labels continue in Figure 5. **c)** BioBloom salt tolerance results at the *mepM. znuA* locus. Results are depicted as in (b). but un-adjusted p-values are used, given that these loci were identified at the genome level using adjusted p-values. Clusters of significant mutations appear to occur within presumed RBS sequences upstream of genes, the *zur* transcription factor binding site (purple). and the *mepM’s* lytM catalytic domain (yellow). **d)** The *ompR. envZ* locus. with results depicted as above. Significant mutations appear to cluster in the presumed regulatory region immediately upstream of *envZ*. and throughout the *ompR* coding sequence, especially surrounding *ompR’s* asp-55 phosphorylation site (purple) and c-terminal domain. **e)** The *hflX. hflK. hflC* locus, with results depicted as above. Significant mutations appear throughout this region, but appear to cluster in the presumed regulatory regions immediately upstream of genes. **f)** The *mraY, murD* locus. with results depicted as above. Significant mutations are detected in upstream and N-terminal regions of these genes. with presumed regulatory effects.

In contrast with rifampicin tolerance which is driven by mutations at a single locus, SNPs across multiple loci appeared to improve salt tolerance. Some of these loci seemed plausibly linked to salt tolerance given existing literature (Figure 4b. green), even if they hadn’t been specifically shown to increase tolerance. We were more skeptical that SNPs in other loci were linked to salt tolerance (Figure 4b. yellow, red). We thus reasoned about the possible mechanisms through which these genes might drive improved growth, and planned to clonally validate a subset of mutations in a salt tolerance assay.

- mraY Significant SNP enrichment occurs at the *mraYlmurD* locus (Figure 4f), both essential genes involved in cell wall biosynthesis and known to be differentially regulated in response to salt stress^44^.
- ompR High-significance SNPs are observed at the *ompRlenvZ* locus (Figure 4d). OmpR is a transcription factor with well-known connection to salt and other stressors, and forms a two-component system with envz^45–47^_ SNPs upstream of envZ are significantly enriched, which may decrease envZ expression. Significant SNPs also occur within ompR’s open reading frame, in many cases clustering around the primary phosphorylation site (Asp55)^48^, and in the C-terminal region where mutagenesis is known to decrease or destroy transcriptional activation^49^.
- **mepM** The highest-significance SNPs identified in this screen occurred in the region surrounding murein endopeptidase M, *mepM* (Figure 4c). *mepM* is one of the core D,D-endopeptidases of *E. coli* involved in cell wall biogenesis, and *mepM* inactivation is known to confer salt sensitivity, specifically linked to its C-terminal Zinc-dependent *LytM* domain^50^. We observed enrichment of SNPs in the ribosome binding site and N-terminal region of *mepM*. which we hypothesize to alter expression. as well as within mepM’s catalytic *LytM* domain. While *mepM* loss of function is known to decrease salt tolerance and is implicated in sensitivity to antibiotic and oxidative stress^50,51^, *mepM* mutants have not been linked to increased salt tolerance before to our knowledge.

Additional significant SNPs are associated with the zinc uptake gene znuA. *mepMs* immediate neighbor in the genome. These SNPs are observed upstream of *znuA* and within its N-terminal region, likely affecting expression. The most significant *znuA* hits overlap the known upstream zur transcription factor binding site. *znuA* SNPs could be acting through *mepM* by altering acquisition of zinc cofactors, these mutations could simply be altering transcription of the downstream *mepM* locus, or *znuA* could be acting independently through some other unclear mechanism.

- **hflXKC** Highly-significant SNPs were detected in the hflXKC operon, despite no direct link to salt tolerance we could discern (Figure 4e). *hflXKC* genes are best known for their role regulating lambda lysogeny^52^, and are not connected to salt tolerance in E. *coli* to our knowledge. We hypothesized that *hflXKC* conferred a fitness advantage due to a defective lambda prophage remaining in this strain due to its construction method^18.21^, and would fail to validate clonally in a salt tolerance assay.

Similarly, we were more skeptical of significant but solitary SNPs. lacking a “cluster” of significant neighbors as seen in the loci above. A significant SNP in the *yidK* putative transport gene and an intergenic SNP between the mrr and *hsdR* restriction system genes are examples of these. and we chose these for validation along with examples from all loci above.

### Characterization Of Salt Tolerance Variants

We were interested to validate whether or not the various SNPs surfaced by BioBloom improve salt tolerance outside of the pooled assay context. We reconstructed SNPs clonally in clean backgrounds (Methods) and measured their growth inhibition in a range of salt concentrations in comparison to the parent strain (Methods, Fig S5a). Mutations in the *mepM* region led to salt IC_50_ values of ~6.4%, about 85% higher than the parent strain’s 3.45% salt IC_50_ (Fig 5a). This validates the observation that mutations in the mepM CDS or upstream region confer significant salt tolerance in E. *coli*, which has not been shown before to our knowledge. Where we hypothesized that *hflXKC* SNPs were some kind of artifact, we were instead surprised to see that SNPs selected from all three of these genes significantly improved salt tolerance. Selected *hflC* and *hflK* SNPs improved salt IC_50_ to 6.53% and 6.38% respectively, and our example *hflX* SNP showed a more modest IC_50_ improvement to 5.26%. The solitary mutations. including those from *yidK* and intergenic between mrr and *hsdR* failed to significantly improve salt tolerance in this assay, as expected. Presumably these false-positive SNPs result from background mutations arising during genome editing. or strong skew in the starting library-both sources of variation we believe can be addressed with future changes to experiment and analysis (see Discussion). SNPs from the *ompR* and *mraY* loci were not successfully constructed in this study. but could be a subject of future work. We were also curious whether measurement of salt IC_50_ might differ after pre-acclimating a strain to salt. Strains pre-cultured in 3% total salt, similar to wild-type IC_50_ did not display meaningfully different salt tolerance to those pre-cultured in standard medium, in this assay (Fig S5b).

**Figure 5:**
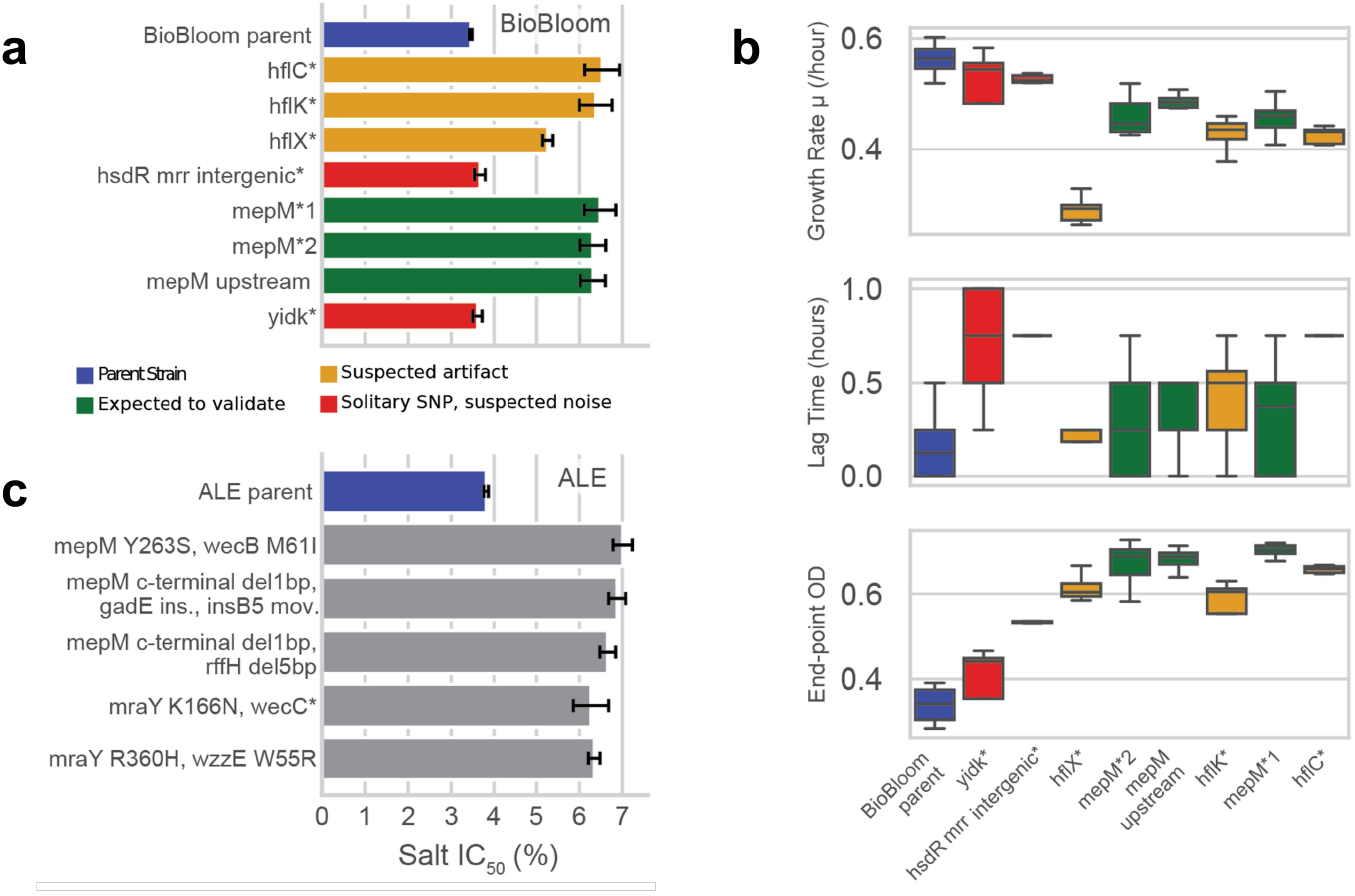
Characterization of salt tolerance variants. **a)** Salt IC_50_ measurement of reconstructed mutations identified using BioBloom. Detailed inhibition curves are provided in Figure S5a. Data is colored by the different classes of mutations chosen from the full-genome data for characterization. Bars depict the mean of replicate measurements and error-bars depict 95% confidence interval of IC_50_ derived from sigmoid curve fitting. b) Growth curve characterization of reconstructed mutations, with exponential growth rate (µ), lag time, and end-point OD depicted. Replicate measurements are summarized by box plots in the style of Tukey. Raw growth curve data is visualized in Figure S6. c) Salt IC_50_ measurement of isolates from ALE. Detailed inhibition curves are provided in Figure S5c. Results are depicted as in (a).

To explore beneficial mutant phenotype in more detail, we next measured growth curves in LB with 5% salt, the condition used for pooled BioBloom selection experiments (Fig S6). Somewhat unexpectedly, mutants with improved salt IC_50_ do not exhibit improvements in either exponential growth rate nor lag time, and instead continue growing to attain a higher terminal biomass yield in these conditions (Figure 5b). The three SNPs in the *hflXKC* operon display significant decrease in exponential growth rate, but still accrue increased biomass in 5% salt, with *hflX”* growing the slowest. *mepM* and *znuA-* associated mutants also show mildly decreased growth rate. but produce the highest terminal OD in these batch growth experiments (Fig 5d).

Different stresses or culture regimes may surface mutants with various growth phenotypes. For example a turbidostat might select for increased growth rate to a greater extent than sequential batch cultures used here-further characterization of selected beneficial mutants can help tune selection conditions to achieve the desired results.

Having confirmed that the SNPs surfaced by BioBloom improve salt tolerance in secondary assays, we were next interested to see how such point mutants compared to multi-mutants surfaced by more traditional adaptive evolution methods. We had previously conducted a one-month serial passaging ALE experiment beginning with *E. coli* MG1655 as a parent in 5% salt (Methods). We isolated 10 clones from the resulting evolved population, full-genome sequenced them to identify their SNPs, and characterized their salt IC50. In contrast with BioBloom variants, strains generated by ALE tend to contain multiple mutations, including movement of mobile elements and possible genome rearrangements. There is overlap between the alleles found in BioBloom and ALE, with both techniques identifying mutations in the *mepM* locus and within *mraY*. The lllC50 improvements generated by BioBloom SNPs and one month of ALE strains were of similar magnitude (Figure 5a,c), with definitive comparison somewhat complicated by a slight difference in the IC50 of the parent strains for the two experiments. BioBloom enabled us to observe a broader diversity of beneficial alleles than those identified by ALE and to confidently attribute phenotypes to specific SNPs, and finds these mutations more quickly.

## DISCUSSION

Here we report BioBloom, a new method for barcoded saturation mutagenesis of bacterial genomes. We constructed libraries containing 99% of possible saturating SNPs in *E. coli*, and used these libraries to identify beneficial mutations under salt and antibiotic selection. BioBloom enables rapid, pooled discovery of beneficial SNPs at nucleotide resolution, and does so genome-wide. In contrast to ALE, BioBloom functions on shorter timeframes, uses smaller working volumes, makes more efficient use of sequencing reads, and ascribes phenotypes to specific mutations without follow-up experiments. The beneficial mutations we identified with BioBloom were more diverse and numerous than those obtained after 30 days of ALE, and BioBloom-identified single mutations achieved similar phenotypic improvement to multi-mutation strains resulting from ALE. In contrast to CRISPRi or TnSeq which tend to ablate the function of genes, BioBloom can augment their function by introducing SNP variants. As a result, we found that BioBloom surfaced catalytic or otherwise critical protein regions and regulatory sequences with no prior knowledge. In addition to highlighting these regions or motifs, BioBloom data is rich enough to suggest strategies for altering them, making it a powerful tool for both understanding and engineering strains for new functions. The speed and efficiency of BioBloom may help identify beneficial SNPs in situations where ALE might be impractical, such as expensive media or growth conditions, very slow-growing strains, or small populations. BioBloom also seems well-suited to identify beneficial SNPs in complex, mixed communities, where multiple organisms complicate the sequencing and isolation steps required for ALE.

Although powerful, BioBloom is not universally preferable to ALE, TnSeq. or other approaches, especially in organisms with limited genetic tools. Synthesis of a complete set of pro-donors (approximately $30k at current prices) remains a substantial barrier. The organism must be amenable to high-efficiency transformation in order to obtain, even, saturating libraries effectively sampling tens of millions of mutations. Most of all, retron recombineering must function efficiently in the target organism; this remains a barrier in many cases, though progress is being made toward efficient retron editing efficiency in diverse strains^**53**^.

These requirements may be relaxed for some studies; targeting a subset of the genome could decrease library size required, and a lower editing efficiency in a new organism would likely still identify beneficial alleles with large effects, given that these edited alleles could purify themselves out of the population during selection^18^. There are many scenarios in which ALE, TnSeq, or other methods remain preferable to BioBloom, nonetheless future work adapting BioBloom to other bacteria and applications holds significant promise.

Further development of BioBloom data analysis could yield additional insights. Analysis of mutation frequencies across multiple timepoints rather than the single-timepoint analysis performed here is expected to improve sensitivity and accuracy. Aggregating signal within regions may reveal cases where single SNPs are not statistically significant, but together are highly significant. an approach commonly used in TnSeq analyses to identify significant regions. SNP effect prediction57 could help filter, prioritize, and interpret beneficial SNPs. SNPs were identified in pro-donors by alignment in this study, but an approach requiring exact match to expected sequences might be more robust in the face of synthesis and sequencing error. Future work is needed to further explore these and other potential improvements in analysis.

Future laboratory work could further improve the BioBloom methods. As with other pooled enrichment assays, BioBloom data quality depends on library evenness, selection strength, population size, and sampling schedule, and needs may differ across different selections. We identified library skew as a critical factor that affects the saturation achieved and the sensitivity of measurements. Decreasing skew could be achieved in simple ways; developing the PCR oligo pool amplification protocol, improving transformation to recover a larger libraries with even better coverage. or passaging larger populations of cells to minimize genetic drift. Improved retron constructs^**23**,**24**,**53**^ or incorporation of known oligo recombineering design principles^**22**,**54**^ could allow for shorter induction period, reducing skew while maintaining or improving editing. We also would have benefitted from earlier sampling during selection. given a small number of highly-beneficial alleles dominated the results of our BioBloom antibiotic resistance experiment and likely obscured less-fit but still significant alleles. We’ve incorporated some of these findings into BioBloom-Ec.-2.0 prior to sharing with others.

### Distribution Of BioBloom Libraries To The Scientific Community

We anticipate that a full-genome SNP library may be useful across diverse projects, akin to shared genome-wide resources like the Keio Collection^58^, the GeCKO Crispr library^59,60^, the human ORFeome^**61**^. and others. Taking into account our findings using BioBloom-Ec.-1.0 in this study, we chose to prepare BioBloom-Ec.-2.0 (Table Sl) prior to distribution, with improvements in experimental feasibility, analysis, and data quality in mind: i) libraries were assembled and transformed in triplicate, to spread any “skewed” representation of mutants across biological replicates as much as possible and help filter results. ii) While BioBloom-Ec.-1.0 was constructed in a base strain with the temperature-induced recombination function at the *tibioAB* locus^21^, in BioBloom-Ec.-2.0, this locus was restored to wild-type, enabling the strain to be selected in minimal media and at 37C. iii) Deep-sequencing was performed to generate baseline pro-donor frequency data for the initial. induced library, sparing this effort and expense for future users and helping enable more sophisticated mutant fitness analysis across multiple timepoints. BioBloom-Ec.-2.0 is available from Addgene as kit <u>1000000273</u>, and baseline frequency data this library is available at <u>doi.org/10.6084/m9.figshare.31053121.v3</u>. Contact for other inquiries.

## METHODS

### Synthesis And Construction Of Full-Genome E.Coli Library

Pro-donor sequences were designed with a custom python script. The script iterates through each base in the genome, obtains the surrounding 82 base-pair genomic window, and replaces the 41st base of the window with an “N”. The sequences are then reverse-complemented depending on which genome replichore they reside in, such that the final ssDNA donor after transcription and reverse-transcription anneals to the lagging strand of replication for optimal editing efficiency^22^. Adapter sequences were added to the 5’ and 3’ end of each pro-donor sequence to allow amplification and assembly by Bsal-golden gate. These pro-donor sequences are designed to insert into the msd region of the Ee 1 retron for retron recombineering. following existing, published work^18.23^. Notably, because Bsal-golden gate was used, the 261 Bsal recognition sequences present in the genome likely obtained depleted coverage. Total Bsal sites present in all output sequences were tracked. These 120bp pro-donor sequences and their golden gate adaptors were synthesized as an oligonucleotide pool by a commercial semiconductor method^26,62^(Genscript). At the time of this writing, this method is more cost-effective than alternative “inkjet printing” methods^63–65^, and is able to accommodate degenerate base mixes.

We also explored an alternative “soft-randomized” approach, in which bases were mutagenized during synthesis by including a percentage “N” such as 1.5%, inspired by related approaches in Oligonucleotide Recombineering^66^ (Fig. S4). This in principle can save substantial synthesis cost because donors can cover larger windows of the genome like 30bp, rather than 1bp. However we found that it resulted in more uneven sampling of mutations than the strategy above where a single position is diversified, and potentially issues with amplification of library members given that amplification sequences were diversified.

Synthetic oligo pools were amplified using Platinum Superfi II polymerase (ThermoFisher Scientific) for a total of 13 cycles in 50 ul reactions. 50 ng of input material was used, following amplification guidelines from the manufacturer^67^. However, we differ from these guidelines by using a single amplification step, given that multiple rounds of amplification were found to be unnecessary. 13 cycles was the minimum required to obtain material for downstream steps, to minimize PCR bias. A linear backbone vector was generated by PCR from the pMS_375 retron recombineering plasmid with chloramphenicol resistance marker^18,68^, a gift from George Church (Addgene plasmid #182132). Both insert and vector fragments were cleaned/concentrated using magnetic DNA-binding beads (Sergi Lab Supplies), and assembled using NEBridge Golden Gate Assembly Mix (Bsal-HF-v2, New England Biolabs (NEB)) following manufacturer recommendations. except that roughly double the recommended DNA per reaction volume was used, the reaction was run for 12 hours total. and three 400 ul reactions were used, to maximize the output CFU following transformation.

After cleaning and concentrating the resulting assemblies they were electroporated into NEB 10-beta electrocompetent *E. coli* (NEB), targeting 0.5-1 ug DNA per 100 ul electroporation in 2mm electroporation cuvettes, using the “Ec3” setting on either the Bio-Rad MicroPulser or GenePulser Xcell instruments. 9 total such electroporations were performed and recovered in Recovery Medium (Intact Genomics) for 1 hour at 3,C. This recovery and all liquid cultures in this work were grown with shaking at 220 rpm. Serial dilutions were then plated to Lysogeny broth (LB) agar medium with 30 ug/ml Chloramphenicol (CM30) to estimate total library CFU. All recovery cultures were pooled and cultured in 50 ml of PlasmidPlus medium (HTS labs) with CM30 for approximately 16 hours at 3,C. Serial dilution plating estimated that each electroporation achieved greater than 1 × 10^8^ CFU, with total pooled CFU estimated at 2.8 × 10^9^, or ~140-fold library coverage across 2 × 10^7^ library members. Plasmids were isolated using the ZymoPure II plasmid Midiprep kit (Zymo). We refer to this plasmid library as “BioBloom-Ec-1.0-plasmid”.

This plasmid library was transformed into the retron editing strain, an *E. coli* derived from MG1655 containing ΔsbcB and ΔrecJ alleles to boost retron recombineering rates^18,30^. Notably, this strain contains a temperature-inducible, defective lambda prophage to facilitate strain construction, and it’s recommended to grow it at 30-32°C^**21**^. This function was removed for construction of BioBloom-Ec-2.0 library. Transformation was carried out by electroporation as described above, using frozen electrocompetent cells prepared commercially (Intact Genomics) from the retron editing strain. Fresh electrocompetent cells prepared in-house can achieve sufficient library scale as well^28^. 9 electroporations were performed as described above, except using these cells and 500 ng plasmid each. Following transformation recovery at 30’C, serial dilution plating was again performed to estimate library yield. Recovery cultures were pooled and diluted into 50 ml total of LB supplemented with both CM30 and 0.4% L-arabinose to induce retron editing. Serial dilution plating estimated that each electroporation achieved >4 × 10^8^ CFU, with total pooled CFU estimated at >1 × 10^10^ CFU, or more than 500-fold library coverage. Following culture at 30’C for 16 hours, 0.5 ml of retron editing cultures (approximately 1.2 × 10^9^ cells or 60-fold library coverage) were again diluted into 50 ml LB+CM30+0.4% L-arabinose to continue editing in triplicate. This was the smallest bottleneck that populations experienced, and would be our first target for improving library skew. These cultures were grown for approximately 20 hours at 30’C for a second round of editing. Induction in this way resulted in ~16 elapsed generations of retron editing, similar to the ~20 generations previously resulting in >50% editing across various alleles^18,23^. We refer to this library as “BioBloom-Ec-1.0-edited” and selection was performed on these cultures.

### Selection Conditions For Full-Genome E. Coli Library

Rifampicin selection was performed by first preparing O-tray dishes (Corning) filled with 200 ml of LB supplemented with 50 mg/ml Rifampicin. 2 ml of cultures from either BioBloom libraries or control conditions were plated to these dishes with plating beads.

Salt selection was performed by diluting 0.5 ml edited libraries into 20 ml of LB medium with an additional 4% w/v Sodium Chloride (salt) added in 50 ml Bio-Reaction tubes (Cole-Parmer) cultured with shaking at 30’C. After 24 hours of growth this dilution was performed again. with ‘timepoint’ representing the number of these 40X dilution/growth cycles performed.

Solid medium selections were harvested using plate scrapers and resuspended in lX PBS buffer before centrifugation at 4000g for 10 minutes. Liquid medium selections were harvested by centrifugation directly. In both cases. OD_600_ was measured in order to pellet approximately 1 × 10^10^ cells, a typical input for a plasmid prep. The resulting pellets were frozen at −20’C and plasmid DNA was later extracted using a Mini-prep protocol (ThermoScientific GeneJet).

### Wild-Type And Mutagenic Control Conditions For Rifampicin Selection

To compare BioBloom to simple mutagenesis, selection, and shotgun sequencing, a Wild-Type culture of E.coli MG1655 was grown for 16 hours at 37°C in 5 ml LB medium, and 2 ml culture was plated to Rifampicin O-trays, in the same manner as above. To explore the effect of mutagenesis in this comparison, the mutagenic plasmid MP6^41^ was first transformed into the same strain, a single transformed colony was grown for 16 hours in 5 ml LB medium with CM30 and 0.2% L-arabinose to induce mutagenesis. MP6 raises the mutation rate by as much as 1 × 10^5^-fold displaying a substantial bias toward transition mutations but a more even mutation spectrum than many mutagenesis methods/genotypes^4l, 69^. These control experiments were performed in quadruplicate.

After 20 hours of growth on solid rifampicin plates at 37°C, colonies were visible on solid medium. Wild-type produced tens of colonies, MP6 cultures produced hundreds, and BioBloom libraries produced a lawn. Material was harvested by scraping as above, except that controls were subjected to DNA extraction (Wizard Kit. Promega). rather than plasmid isolation. Genomic DNA was “tagmented” using established methods (Nextera-XT. lllumina), and indexed for sequencing by amplification using published Nextera indexes^70^. This produced shotgun genomic libraries, in contrast with the targeted amplicon libraries used for BioBloom.

### Amplicon Sequencing Of BioBloom Libraries

Extracted plasmids were first amplified using the “fwd” and “rev” primer sets (Table S3).Forward and reverse primers were mixtures incorporating degenerate bases to increase amplicon diversity at the beginning of lllumina read l and read 2. These reactions were performed with 05 DNA Polymerase Mix (NEB) in 40 µL reactions using 2 µL plasmid DNA as template, annealing at 65°C for 19 cycles. Product was treated with l µL DNA Exonuclease I (NEB) at 37°C for 10 minutes followed by heat inactivation at 80°C for 20 minutes. l µL of this treated material was used as a template in a second indexing PCR using dual indexing primers (Zymo-Seq UDI primer sets, Zymo research), a 20 µL reaction annealing at 66°C for 5 cycles only. The resulting product was purified using magnetic DNA binding beads (Sergi lab supplies) and pooled for sequencing.Sequencing was performed alongside other genomic and amplicon samples using a 2x-l50 bp protocol on a NovaSeq instrument (Seqcenter).

### Analysis Of Sequencing Data

All sequencing analysis is available in iPython notebook format at github.com/Pioneer-Research-Labs/BioBloom_preprint, and data is accessible at 10.6084/m9.figshare.31053121.

Targeted amplicon sequencing of BioBloom samples yielded ~11-19 × 10^6^ read pairs.After merging with fastp^71^, merged reads were trimmed to donor sequences using cutadapt^72^. Notably, donors were covered by both Rl and R2. and fastp performs error correction across all merged bases, effectively lowering the sequencing error of these regions^71^. Identical sequences were identified and counted using a custom bash command, discarding any sequences appearing fewer than 10 times, which are unlikely to be under selection. The resulting de-duplicated sequences were aligned to references using bwa -mem as above. Where library coverage at genomic positions was determined. alignment files were summarized using samtools coverage. Custom scripts parsed alignment files for mutations while logging information about mutation position within a donor sequence.Mutations were filtered for those that occurred at the expected “N” editing position at donor base 41, or in the two surrounding bases to accommodate for variation in read-pair length.We found that this captured the majority of intended mutations. While depressing the signal from sequencing errors. Notably, unintended mutations resulting from synthesis do occur at other positions and produce plausible effects which may be of interest-we detected a substantial number of single base-pair deletion mutations, not intended but likely resulting from synthesis, many of which produced plausible effects.

p-values for mutations were obtained using a permutation test; wherein a mutation’s count is compared to a full, scrambled set of unlabeled mutation counts within the same sample, to infer the likelihood of a mutation count at least that extreme. p-values were summarized across replicates using Stouffer’s method^73,74^. P-values were adjusted for multiple hypothesis testing using Benjamini-hochberg correction. Results were visualized using python packages matplotlib, seaborn, plotly express, and venn2.

Control conditions which underwent shotgun genomic sequencing received ~18-27 × 10^6^ read pairs. Reads were trimmed and merged using fastp^71^, and the resulting merged and un-merged files were independently aligned to the U00096.3 MG1655 reference^75^ using bwa -mem^76^ and the resulting .barn files combined using samtools merge^77^. These alignments were then summarized by the variant caller lofreq^78^, which is specialized for low-frequency, population-level variant analysis. After inspection, variants with Allelic Fraction (AF) greater than 98% were judged to be differences between the starting strain and reference genome, and were filtered from analysis. Variants were analyzed using statistical methods detailed above for BioBloom. but using the total number of observations of a given mutation in shotgun sequencing as the input data.Using the Allelic Fraction of mutations as input data is an alternative method, and was found to produce largely identical results.

### Re-Construction And Investigation Of Putative Salt-Tolerance Mutations

A set of 13 mutants showing significant p-value in salt were chosen for validation, including both members suspected to be true hits *(mepM, ompR, mraY* alleles), those suspected to be artifacts *(hflXKC, hsdR* alleles), and others. For each allele, a short 77 basepair oligo recombineering donor was designed using MODEST^79^, and primers to incorporate these donors into retron constructs were designed. The retron vector pMS_375^18,68^ was amplified with these primer pairs and assembled by KLD enzyme mix (NEB).Clonal plasmids were isolated and their donor region verified by sanger sequencing.*hflXKC* is annotated as a regulator of prophages including lambda. To prevent phenotypic effects acting through the lambda prophage rather than salt tolerance, we first removed the lambda prophage in this strain, restoring the *bioAB* locus in our test strain using established methods^21,58^. Retron plasmids were introduced into these strains by electroporation. Editing was induced with arabinose and CM30 selection as described above but in l ml volumes. Colonies were isolated by streaking. and mutant alleles were confirmed by sanger sequencing of edited loci. 8 mutations were successfully isolated and are investigated in this work. These isolates were grown for 16 hours LB medium in the absence of CM30 and with 400 µg/ml novobiocin to encourage plasmid curing^80,81^, and streaked for single colonies.Colonies were screened for loss of growth on CM30 (>90% of colonies) and alleles further confirmed by whole genome sequencing (Plasmidsaurus).

Salt inhibition was measured by first pre-culturing strains in LB medium for 16 hours, then diluting 100-fold into l ml cultures of LB medium with varying concentration of added salt and culturing an additional 24 hours at 37°C. Optical density (OD of these cultures was read on a plate reader (Spectramax 340pc) after diluting 3-fold into a 2% salt solution. IC was determined by a sigmoid fit to these data using a python script. Growth phenotypes in salt were further investigated by monitoring growth curves in 4% salt over time in a shaking, incubating plate reader (BioTek Synergy HT) at 37°C.OD_600_ values from this experiment were analyzed using the croissance^82^ package in python.

### Modeling Of Sampling And Mutations

To estimate the fraction of all SNPs sampled by a BioBloom Library, we began with a simple Coupon Collector model^83^ in which the fraction of all mutations sampled equals 1 - ((1 - 1 / S_m_) ** N_m_), where S_m_ is the size of the SNP mutation space (1.5 × 10^6^ for a genome of 5 × 10^6^ basepairs, and N_m_ is the number of mutations drawn. This model was extended to account for ~25% of library members being Wild-Type sequence and thus contributing O mutations, and was further extended to estimate the effect of drawing from a skewed distribution. The distribution of library members was modeled as log-normal with a mean normalized to 1.

To estimate the fraction of all mutations sampled in a bacterial culture, a simple model was constructed in which the total number of mutations is the number of divisions * mutation rate per basepair (1 × 10^−10^ mutations/ basepair^69^) * genome length (~5 × 10^6^ basepairs). The number of divisions in a culture was estimated as the number of cells in that culture, following oft-referenced work^33^. The Coupon Collector model was once again used to estimate the fraction of mutations sampled when randomly drawing mutations in this manner. This model was extended to partially account for mutation spectra, namely the lower rate of transversion mutations (2 × 10-11 mutations/basepair^69^) and assuming that transversions are approximately½ of total possible SNPs. The number of cells in a confluent culture was estimated as 2 × 10^9^/ml, and the total number of cell divisions in a 30X serial batch culture is estimated as 30X the cell divisions represented in a single batch culture. This model was further extended to account for editing efficiency when determining the population size of edited cultures required for selection experiments (Fig S2b).

### Adaptive Laboratory Evolution, ALE

*E. coli* MG1655 was grown in LB liquid medium (Hi-media, “Miller” formulation) with an additional 4% weight/volume sodium chloride added, for a total sodium chloride content of 5%. Cells were grown across 3 replicates in 1.05 ml cultures in a 96-well deep-well plate with shaking at 1000 RPM at 37°C. 50 ul of cells were transferred into fresh media approximately every 24 hours, for one month. Halfway through the experiment, we chose to double the number of replicates by splitting each well into two for a total of six wells, starting after 16 days of culture. OD was measured at dilution time to monitor growth and anticipate potential problems. Cultures were cryo-preserved in 15% glycerol at −80°C every week. Final cultures were streaked out from the terminal culture after 30 days. 10 isolates obtained from the streaks were subjected to full-genome sequencing (Angstrom Innovation), and salt inhibition testing by IC_50_. as above.

## Acknowledgements

This research was funded by The Astera Institute, the Experiment Foundation, The Align Foundation, and Schmidt Sciences. We thank Gemma Alderton, Michaela Hinks, Samantha Piszkiewicz, Shantal Al Habib, Kayla Young, Amanda Apel, and Akos Nyerges for their comments on the manuscript. We thank Rachel Sevey and Olesia Bushkova for their assistance with scientific communication. We thank Seth Shipman for their early input on the project.

## Open Peer Review

To read existing peer reviews or post your own review, visit either ResearchHub https://www.researchhub.com/paper/11079221 or PreReview https://prereview.org/preprints/doi-10.64898-2026.02.04.703572.

