## Supplemental materials and figures for "BioBloom, a method for barcoded saturation mutagenesis of an entire bacterial genome"

### GLOSSARY

**ALE, Adaptive Laboratory Evolution.**
An experimental approach in which populations are cultured over many generations under defined selection conditions to enrich beneficial mutations.

**CDS, Coding DNA Sequence.**
The portion of a gene's DNA sequence that is translated into protein, from start codon to stop codon.

**CFU, Colony Forming Unit.**
A unit estimating the number of viable microorganisms in a sample, defined by the ability to form a visible colony on solid medium.

**CRISPR (Clustered Regularly Interspaced Short Palindromic Repeats).**
A genome editing system derived from bacterial adaptive immunity that uses a guide RNA and a CRISPR-associated (Cas) nuclease to target and cleave specific nucleic acid sequences.

**Degenerate nucleotide.**
A nucleotide position in a DNA sequence specified to represent multiple possible bases (e.g., "N" meaning A/C/G/T). For example, a degenerate position in a synthesized oligo can be achieved by flowing in all four dNTPs when synthesizing that base.

**DMS, Deep Mutational Scanning.**
An experimental approach that combines saturation mutagenesis of a defined gene, genomic region, or genome with high-throughput quantitative measurement to assess the functional or fitness effects of individual mutations in parallel.

**IC50, Inhibitory Concentration 50%.**
The concentration of a substance (e.g., drug, salt, inhibitor) that reduces a measured response (commonly growth or activity) by 50% relative to an untreated control.

**RBS, Ribosome Binding Site.**
A sequence in mRNA that recruits the ribosome to initiate translation; in bacteria this typically includes the Shine—Dalgarno sequence upstream of the start codon.

**Retron.**
A bacterial genetic element that encodes a reverse transcriptase and produces a characteristic single-stranded DNA via reverse transcription.

**NGS, Next Generation Sequencing.**
High-throughput DNA/RNA sequencing technologies that enable parallel sequencing of millions of fragments in a single run. Here, we specifically use NGS to refer to short-read sequencing.

**MG1655.**
A widely used laboratory strain of *Escherichia coli* K-12, commonly used as a reference strain in genetics and molecular biology.

**OD, Optical Density.**
A measure of light attenuation by a sample, commonly used as a proxy for cell density in microbial cultures (often measured at 600 nm as OD600).

**ORF, Open Reading Frame.**
A continuous stretch of nucleotides that can be translated into a peptide, beginning with a start codon and ending at a stop codon.

**Pooled Assay.**
An assay that measures the effects of many genetic variants or perturbations simultaneously in a mixed population. Sometimes referred to as a "multiplex" assay.

**Pro-donor.**
A designed nucleic acid sequence that both serves as the template for generating a single-stranded donor DNA used to introduce a programmed mutation into the genome and functions as a molecular barcode that can be sequenced to quantify the abundance of that mutation in a population.

**PAM, Protospacer Adjacent Motif.**
A short DNA sequence adjacent to a CRISPR target site that is required for recognition and cleavage by many Cas nucleases.

**Saturation mutagenesis.**
A mutagenesis strategy that systematically generates all possible mutations at a target site or across a target sequence.

**SNP, Single Nucleotide Polymorphism.**
A variation at a single nucleotide position in a DNA sequence among individuals or genomes.

### SUPPLEMENTAL MATERIALS

#### Table S1: Summary Of BioBloom Libraries Created

| | BioBloom-Ec.-1.0 | BioBloom-Ec.-2.0 |
|---|---|---|
| Description | Full-genome saturation library of *E. coli* MG1655 | Full-genome saturation library of *E. coli* MG1655, cloned in biological triplicate |
| Number of pro-donors | $4.6 \times 10^6$ degenerate pro-donors | $4.6 \times 10^6$ degenerate pro-donors |
| Theoretical library size | $1.8 \times 10^7$<br>(All SNPs, plus 25% wildtype from degenerate oligonucleotide synthesis approach) | $1.8 \times 10^7$<br>(All SNPs, plus 25% wildtype from degenerate oligonucleotide synthesis approach) |
| Number of transformants (plating, CFU), plasmid library | $2.8 \times 10^9$ | Replicate 1: $2.74 \times 10^9$<br>Replicate 2: $3.07 \times 10^9$<br>Replicate 3: $5.41 \times 10^9$ |
| Number of transformants (plating, CFU), editing library | $>1 \times 10^{10}$ | Replicate 1: $4.67 \times 10^9$<br>Replicate 2: $3.93 \times 10^9$<br>Replicate 3: $5.41 \times 10^9$ |
| Computed Skew, **σ** | **σ** = ~1.3 | **σ** = ~0.869 (rep 1), ~0.757 (Rep 2), ~0.900 (Rep 3) |
| Computed Saturation % | >99% | >99% |
| Notes | - | Assembled and shuttled into the final strain in biological triplicate. Available on Addgene, kit #1000000273. |

#### Table S2: Comparison Of Barcoded Microbial SNP-Editing

| | Mechanism | Organism | What It Is | Largest Library Size Achieved | Saturation Percent Of Genome |
|---|---|---|---|---|---|
| BioBloom (this study) | Retron Library Recombineering (RLR) | *E. coli* | Retron-based genome-scale saturation mutagenesis | $\sim 14.8 \times 10^6$<br>(99% of 15M member set) | >99% synthesized, ~>99% observed after editing |
| CREATE[1] | CRISPR | *E. coli* | CRISPR-based saturation editing of two genes, folA and acrB. Reconstruction of targeted mutations across seven genes. | 50,000 | ~0.33% |
| MAGESTIC[2] | CRISPR | *S. cerevisiae* | CRISPR based saturation editing of one gene, SEC14 | 35,000 | ~0.23% |
| Inscripta- Onyx[3] (similar to CREATE) | CRISPR | *E. coli* | CRISPR based saturation libraries of three proteins, FabZ, LpxC and MurA | 17,000 | ~0.11% |
| CRISPEY[4] | CRISPR | *S. cerevisiae* | CRISPR-based | 16,000 | ~0.11% |

**Table S3: Oligonucleotide Sequences**

| Primer Name | Sequence | Description | Tm |
|---|---|---|---|
| oBB_0120 | CTGAGGCAGGTCTCCTTCC | full_genome_amp fwd | 67 |
| oBB_0121 | AGTCCGTCGGTCTCGATTC | full_genome_amp rev | 67 |
| oBB_0122 | GCCACTCGGGTCTCATTCC | upsteam_lib_amp fwd | 69 |
| oBB_0123 | CTGGGCCAGGTCTCCATTC | upstream_lib_amp_rev | 69 |
| fwd_1 | cctacacgacgctcttccgatctTCTGAGTTACTGTCTGTTTTCCTG | retron donor | 65 |
| fwd_2 | cctacacgacgctcttccgatctVTCTGAGTTACTGTCTGTTTTCCTG | retron donor | 65 |
| fwd_3 | cctacacgacgctcttccgatctNRTCTGAGTTACTGTCTGTTTTCCTG | retron donor | 65 |
| fwd_4 | cctacacgacgctcttccgatctNNRTCTGAGTTACTGTCTGTTTTCCTG | retron donor | 65 |
| rev_1 | gagttcagacgtgtgctcttccgatctGTCAGAAAAAACGGGTTTCCTG | retron donor | 65 |
| rev_2 | gagttcagacgtgtgctcttccgatctHGTCAGAAAAAACGGGTTTCCTG | retron donor | 65 |
| rev_3 | gagttcagacgtgtgctcttccgatctNMGTCAGAAAAAACGGGTTTCCTG | retron donor | 65 |
| rev_4 | gagttcagacgtgtgctcttccgatctNNaGTCAGAAAAAACGGGTTTCCTG | retron donor | 65 |

A note on primers: “fwd” and “rev” primers here include degenerate sequences to increase diversity and help improve sequencing quality on Illumina platforms. Notably, the effective diversity observed in sequencing data is still too low to sequence these amplicons alone and we recommend supplementing with 20+% PhiX or running alongside unrelated samples. Melting temperatures were calculated using the NEB “Q5” melt temperature calculator.

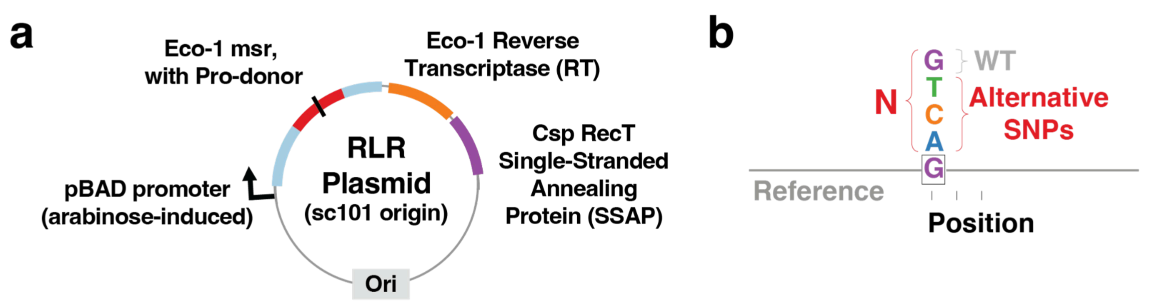


### Supplemental Figure 1

S1a) Additional details of pMS_375 retron plasmid. Arabinose-induced pBAD promoter is indicated with a black arrow. The Eco1 retron with pro-donor sequence is indicated with a red sector, and specific mutation indicated with a black line. Reverse-transcriptase (RT, orange) and Single-stranded annealing protein (SSAP, purple) are indicated, along with a sc101 origin (ori, grey).

S1b) Additional detail on saturating SNPs and degenerate bases. The example genome position indicated here "G" (purple) along a reference genome (grey line) has three alternative bases, T, C, A. The degenerate "N" base contains all 4 G, T, C, A bases, specifying these alternatives, alongside wild-type (WT). This is the conceptual basis for determining the size of a saturating SNP library, the manner in which "N" can substitute for all alternatives, and how 25% of the resulting library is wild-type sequences.

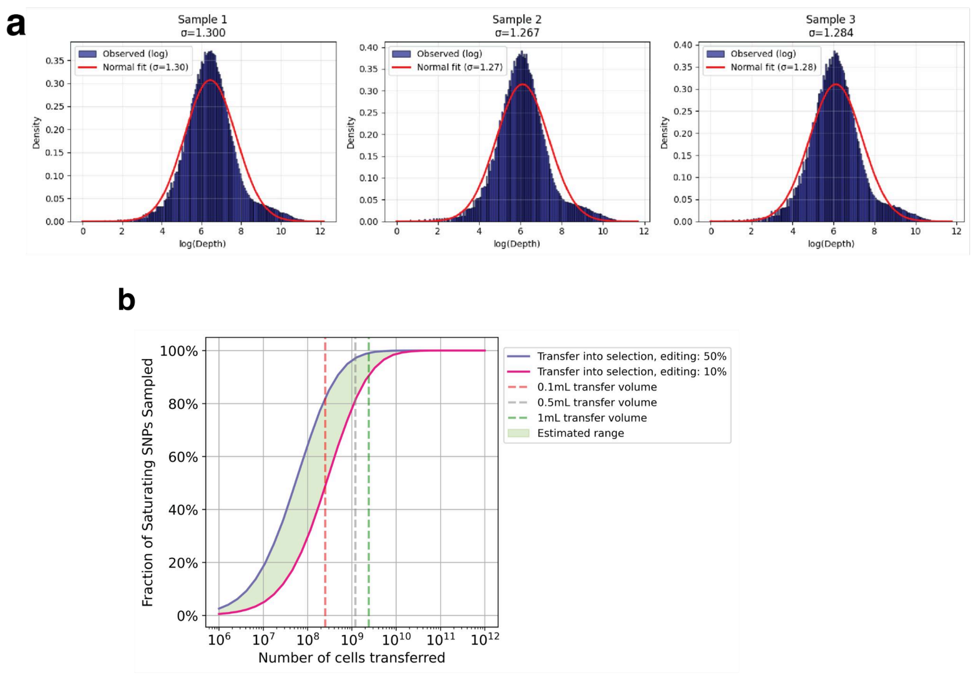


### Supplemental Figure 2

S2a) Distribution of genome coverage at all bases, for all three replicates after induction of editing. A histogram of observed data is shown in blue, alongside a simulated log-normal distribution (red) with the same estimated mean and standard deviation.

S2b) Modeling to determine the volume of edited library needed to perform selection experiments. A coupon collector model is used to model the fraction of saturating mutations sampled from a full set of mutations distributed with **σ** = 1.3, also taking into account different editing efficiency of retrons. If editing efficiency is estimated at 50% (purple) a smaller number of cells are needed to sample the same fraction of saturation as if editing is 10% (pink). We expect that our process produces editing somewhere between these two (Green), and show the range of saturation expected when transferring 0.1mL, 0.5mL, and 1mL of confluent culture.

### SUPPLEMENTAL FIGURES

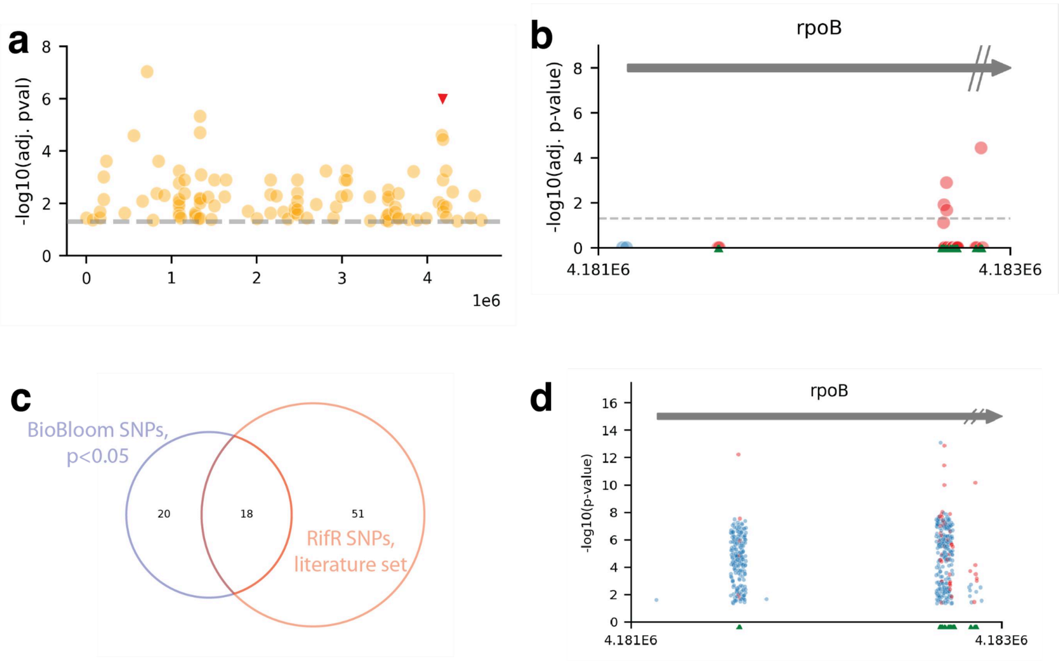


#### Supplementary Figure 3: a companion to Figure 3

S3a) Genome-wide results of rifampicin resistance experiment as depicted in Figure 3e, but showing results for mutagenic populations, bearing the MP6 mutagenesis plasmid[5].

S3b) Results of the rifampicin resistance experiment at the *rpoB* locus, depicted as in Figure 3f, but showing results for mutagenic populations, bearing the MP6 mutagenesis plasmid[5].

S3c) Venn diagram comparing the set of *rpoB* SNPs identified by BioBloom as likely beneficial with p<0.05 (purple), and the set of rifampicin-resistance SNPs expected given a literature summary of such alleles[6] (red). Number of SNPs in different categories is shown.

S3d) Results of the BioBloom rifampicin resistance experiment at the *rpoB* locus, as depicted in Figure 3c except without filtering specifically for mutations at the intended "N" edit position. Far more mutations are detected, but we believe that many of these are sequencing errors resulting from high-coverage sequencing of very successful mutations. All other data depicted in this manuscript uses this filtering method for this reason.

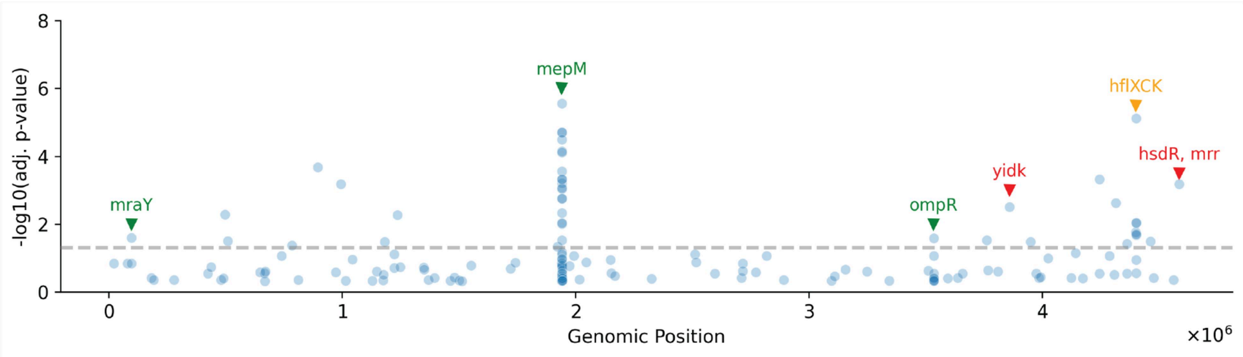


#### Supplemental Figure 4

Results of BioBloom whole-genome salt selection after 3 subcultures in salt, rather than 2 as explored in Figure 3.

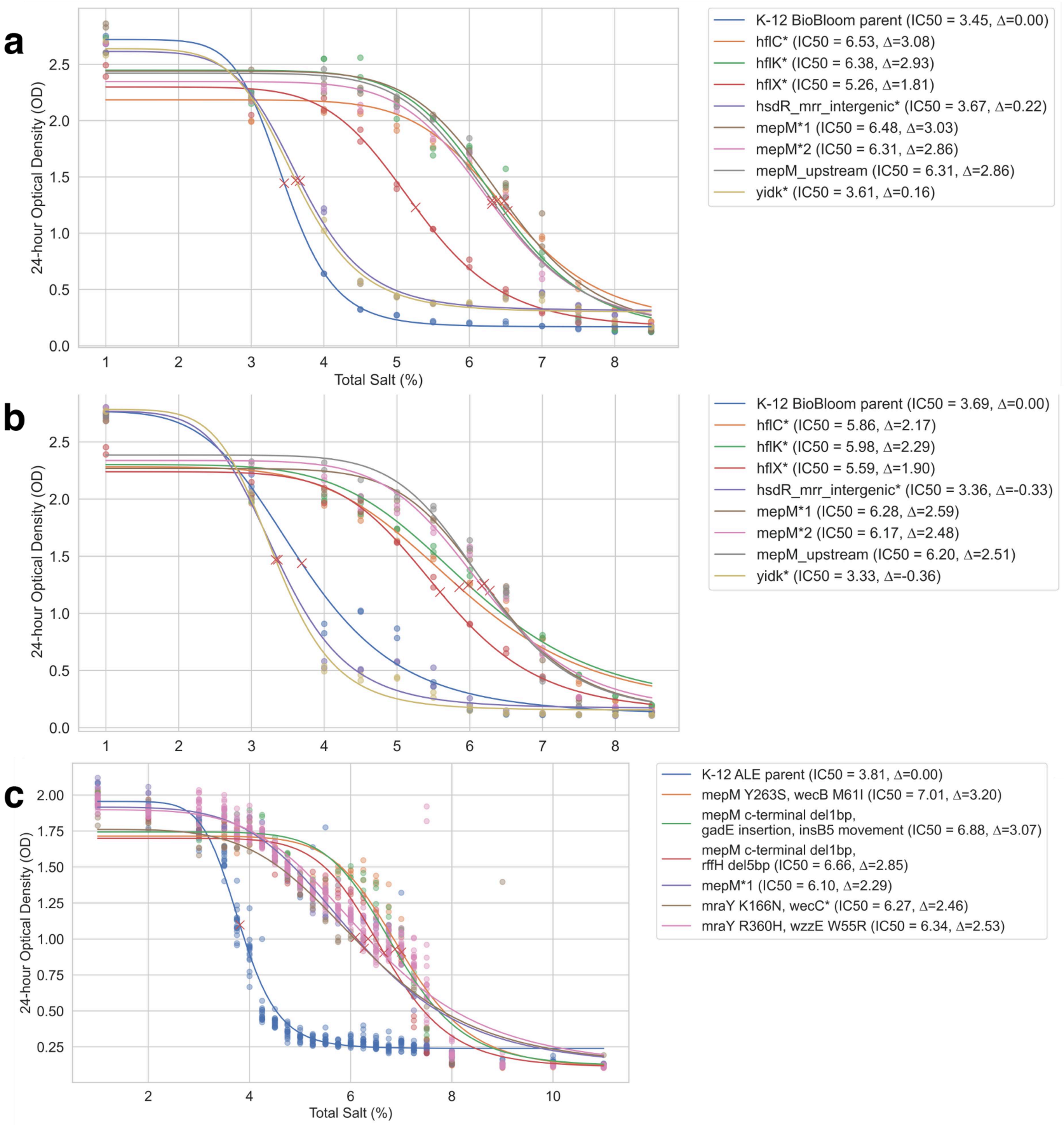


**Supplemental Figure 5: Inhibition curves of selected mutant strains in salt**

S5a) Inhibition curve of BioBloom-identified mutant strains across a range of salt concentrations. $OD_{600}$ after 24 hours of growth (y-axis) is measured across a range of salt concentrations (x-axis). Sigmoid curves are fit to the raw data and used to derive $IC_{50}$ and ΔIC values for mutants, which are shown in the legend.

S5b) Inhibition curves as above, but after pre-acclimation in 3% total salt. Pre-acclimation was judged to be of little importance here.

S5c) Inhibition curves as above, but for mutants isolated from 30 days of ALE.

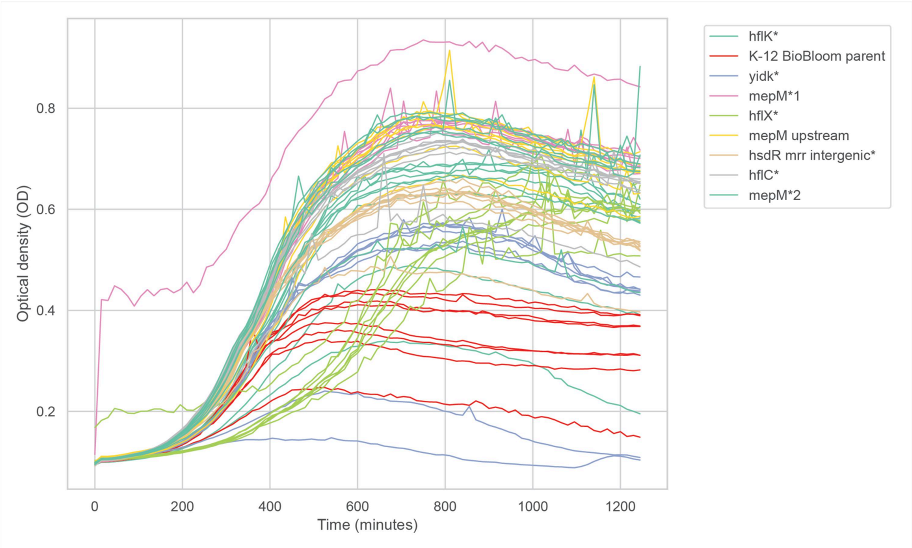


#### Supplementary Figure 6

raw data from growth curves in 5% total salt. Optical density is measured over time for a selection of BioBloom-identified mutants, in replicate. These growth data are summarized in Figure 5b.

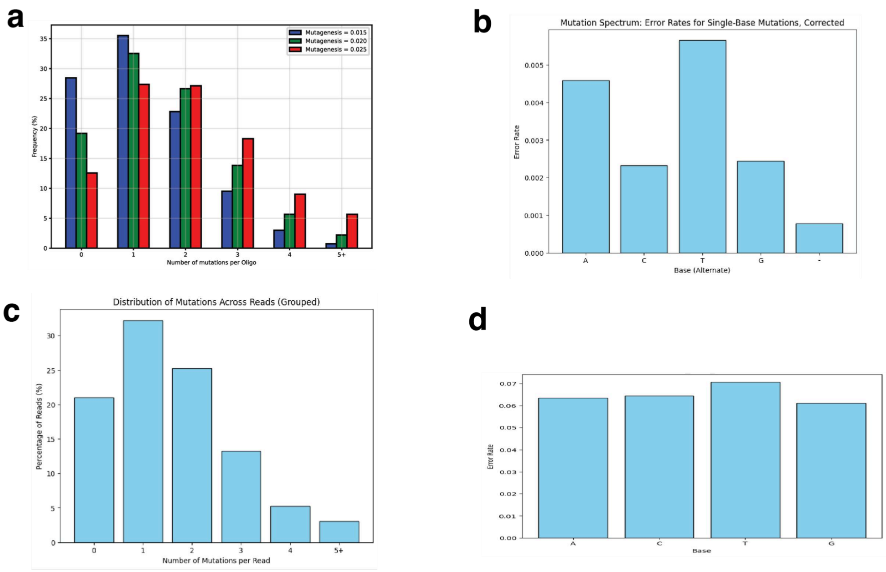


#### Supplementary Figure 7: Early exploration of soft-randomized mutagenesis

S7a) Model of soft-randomized mutagenesis, in which a certain percentage “N” is doped across all synthesized positions of an oligonucleotide[7]. The fraction of oligos with different numbers of mutations are depicted. 2% Mutagenesis was chosen for further testing.

S7b) When oligos synthesized with 2% N were aligned to the genome, the substitution rate for each base was determined. Sequencing error was removed, by subtracting rates obtained in a sample where no mutagenesis was used. This analysis determined that T and A mutations were over-represented when a 3:3:2:2::A:C:T:G mixture was used.

S7c) Actual data obtained when sequencing cloned pools constructed with oligos soft-randomized at 1.5% N. A close match to modeling results was obtained.

S7d) Substitution rates were simulated after correcting the “N” base mixture further, to 0.25:0.3:0.12:0.32::A:C:T:G . This mixture was used as “N” when synthesizing full-genome libraries.
